# Sex-specific retinal anomalies induced by chronic social stress in mice

**DOI:** 10.1101/2020.09.28.317230

**Authors:** E Arsenault, AA Lavigne, S Mansouri, K Francis, TP Bittar, F Quessy, K Abdallah, A Barbeau, M Hébert, B Labonté

## Abstract

Major depressive disorder (MDD) is one of the most common consequences of chronic stress. Still, there is currently no reliable biomarker to detect individuals at risk to develop MDD. Recently, the retina emerged as an effective way to approach the brain and investigate psychiatric disorders with the use of the electroretinogram (ERG). In this study, cones and rods ERGs were performed in male and female mice before and after chronic social defeat stress (CSDS). Mice were then divided as susceptible or resilient to stress. Significant results were only observed in rods ERGs. In males, susceptible mice showed prolonged a-wave implicit times at baseline that were shortened after CSDS. The a-wave was also decreased in both susceptible and resilient male mice after CSDS. In females, rod a-waves were shorter in susceptible than in control mice after CSDS resulting from the latter demonstrating delayed a-waves. Baseline ERGs were able to predict – to some extent – the expression of susceptibility and resilience before stress exposition in male and female mice. Overall, our findings suggest that retinal activity is a presumptive biomarker of stress response and that the ERG could potentially serve as a predicting tool of the stress response in mice.

---

Affecting more than 300 million people worldwide^1^, major depressive disorder (MDD) is a leading cause of disability, imposing a major burden on modern societies^1,2^. Only in United States, more than 14-16% of Americans will require treatment for MDD at one point in their life^3^ and only 20-40% of them will fully recover^4-6^. Noticeably, epidemiologic studies reported that while both males and females are affected, MDD is a sexually dimorphic disease. For instance, women are 1.5-2 times more likely to develop MDD than males^7,8^. Women often exhibit more severe scores of depression associated with higher prevalence of atypical symptoms and co-morbid anxiety^7,9,10^. Additionally, it has been suggested that men and women respond differently to antidepressant treatment^11,12^. While this sexual dimorphism is well recognized, the mechanisms underlying its etiology remain poorly understood.

It is well accepted that disease chronicity worsens patient prognostic. For instance, recurrence after the first MDD episode is up to 30-60% and it increases with each subsequent episode^13,14^. Moreover, the longer the depressive symptoms are present, the harder they are to treat^15,16^. Therefore, early diagnostic is critical to decrease the long-term prevalence of MDD. However, there are still very few tools and biomarkers available to help clinicians. Studies investigating the immune system in MDD patients reported various changes associated with the disease^17,18^. Variations in serum levels of the brain-derived neurotrophic factor (BDNF) have also been associated with response to different antidepressants^19,20^ and changes in the activity of the HPA axis^21,22^, including circulating cortisol, are hallmark alterations consistently associated with the expression of the disease^23^. The cellular complexity found in blood, along with the heterogeneity and severity of the disease, interferes with the reproducibility of these findings, especially during its prodromal phase. Consequently, there is still no reliable biomarker allowing the early detection of individuals at risk to develop MDD.

Alternatively, electroretinography (ERG) has emerged as a non-invasive and reliable approach to investigate psychiatric disorders^24,25^ given that during the embryonic phase, the retina develops from the same cells as the brain^26^. More specifically, the eye development is triggered during early embryonic phases by the activity of the master control gene PAX6 that sequentially induces the outgrowth of the optic grooves, transforming into optic vesicles following the closure of the neural tube and ultimately developing into optic cups^27^. The inner layer of the optic cup, composed of neuroectoderm, gives rise to the retina^27^. The ERG is a biopotential signal generated by the retina in response to a flash of light. Both rod (night vision) and cone (day vision) responses can be isolated and quantified depending on the state of retinal adaptation to darkness or light, respectively^28^. A typical ERG waveform is composed of a negative component called the a-wave, originating from photoreceptors, followed by a larger positive component called the b-wave produced by bipolar cells^28^.

Interestingly, abnormal retinal activity has been consistently reported in many psychiatric disorders including schizophrenia (SZ), bipolar disorder (BP), children at high risk for psychiatric disorders, seasonal affective disorder (SAD), and MDD^29-32^. For instance, prolonged cone b-wave implicit time (time from onset of flash stimulus to the peak of the wave) and reduced mixed rods/cones a- and b-wave peak amplitudes have been reported in MDD patients^33^. Interestingly, the cone b-wave delay identified in MDD patients was similar to the anomalies found in SZ^34^ and BP patients^32^ whereas MDD mixed-rod-cone alterations were common to BP, SZ and children at high risk to develop a psychiatric disorder^31^. MDD patients who responded to antidepressant treatment (Duloxetine) were also shown to exhibit higher rod b-wave amplitude than non-responders before treatment^30^ suggesting that ERG could be used to predict treatment response. Sexual dysmorphism was also examined revealing lower mixed rods/cones b-wave amplitudes in females suffering from SAD compared to healthy female controls with opposite effects in males with SAD^35^ suggesting that ERG may be sensitive to sexual dimorphic features in psychiatric conditions. Finally, alterations have also been reported in different mouse models of depressive and psychosis-like behaviors^36^. For instance, decreased rod retinal sensitivity was reported in DAT-KO mice^37^ and cone b-wave implicit time was prolonged in Tph2-KI mice^37^ suggesting that ERG signals in mice can provide interesting insights into the etiological mechanisms underlying stress response in males and females.

In this study, we tested whether chronic social stress impacts retinal activity in male and female mice and reproduces the retinal alterations observed in MDD patients. To do so we used a model of chronic social defeat stress (CSDS) known to reproduce behavioral features relevant to the clinical manifestations of MDD in men and women^38^. CSDS involves subjecting mice to repeated bouts of social subordination, after which a subset of mice exhibit depressive-like behaviors defined by anhedonia, anxiety-like responses and social withdrawal^39,40^. Interestingly, CSDS in males and females also allows differentiating a subpopulation of resilient mice resistant to the anhedonia and social deficits seen in susceptible mice^41-43^. We used ERG to test whether changes in rod and cone activity associate with the expression of susceptibility and resilience to CSDS in males and females and whether ERG signals before stress exposure could predict stress responses in both sexes.

## Material and Methods

### Animal

C57bl6 male (*n* = 49) and female (*n* = 43) mice aged 7-8 weeks old and CD1 males aged 4-6 months old were obtained from Charles River Laboratories (USA). Mice were fed *ad libitum* at 22–25°C on a 12-hour light/dark cycle. CD-1 mice were singly housed except during social defeats. All other mice were housed in groups (4/cage) before and singly housed after social defeat episodes. In total, 35 males and 30 females were subjected to CSDS. Fourteen males and 13 females served as controls. All experimental procedures were approved by Université Laval’s Institutional Animal Care Committee in respect with the Canadian Council on Animal Care guidelines.

### Chronic Social Defeat Stress (CSDS)

CSDS was performed in males^39^ and females^40^ as described before. Briefly, in males, C57bl6 mice are first introduced to unknown aggressive retired CD-1 breeders in their cages for a period of 5 minutes during which C57bl6 are attacked by the resident mice. Following this initial phase, C57bl6 mice are moved to the other side of the divider for 24 hours allowing a continuous sensory contact with the CD-1 mouse without physical harm. The same procedure is repeated over 10 days with new unknown CD-1 mice every day.

A very similar approach is used in female with the difference that the base of the tail and the pubis of the C57bl6 female mice were soaked with 50µl of male urine before each defeat bouts. All C57bl6 female mice were associated with a different male urine so that CD-1 mice never encountered the same urine twice. Urine collection was performed 1 week preceding social defeat stress with metabolic cages (Tecniplast Group, Tecniplast Canada, Canada). Male mice were housed in metabolic cages overnight. Urine was collected each morning and stored at 4°C after being filtered. The 5-minute social stress bouts were interrupted every time CD-1 male mounted the C57bl6 female mice. Both male and female controls were housed 2/cage separated by a perforated Plexiglas® acrylic divider and housed in the same room as the mice undergoing CSDS.

Social-avoidance behavior was assessed with the social interaction test (SI) 24 hours after the end of the social defeat paradigm as described before^39,40^. Briefly, SI test consisted of two phases of 150 seconds each. In the first phase, C57bl6 mice explored the arena with no target CD-1 aggressor in the social interaction zone. This initial phase was followed by a second exploratory phase but this time in the presence of an unknown target CD-1 aggressor maintained into a mesh-wired enclosure within the social interaction zone. Time spent in the different zones of the arena was automatically recorded through ANY-Maze 4.99 using a top-view camera (ANY-Maze Video Tracking Software, Stoelting Co., USA). Based on social interaction ratios (time in interaction zone with social target/time in interaction zone without social target), defeated mice were designated as susceptible or resilient: susceptible ratio < 1.0; resilient ratio > 1.0. This measure of susceptibility versus resilience has been shown to correlate with other defeat-induced behavioral abnormalities such as anhedonia (e.g. decrease in sucrose preference) and an increased sensitivity to inescapable stresses^41-43^.

### ERG Recordings

The ERG procedure and equipment were the same as the ones used by Lavoie, et al. ^37^, except for the luminance stimuli. All animals were anaesthetized with an intraperitoneal injection of a Ketamine (80 mg/Kg) and Xylazine (10 mg/Kg) mixture under a dim red light. The left eye cornea was anesthetized with 0.5% proparacaine hydrochloride (Alcaine, Alcon Canada, Canada) and the pupil was dilated with 1% tropicamide (Mydriacyl, Alcon Canada, Canada) 10 minutes prior to testing. Lubricant eye drops (Systane Gel Drops, Alcon Canada, Canada) were used to prevent dryness of the cornea of both eyes. A loop shaped DTL electrode (Shieldex 33/9 Thread, Statex, Germany) was placed directly on the left eye cornea to record ERG signals. Reference and ground subcutaneous electrodes (Grass Technologies, Astro-Med, Canada) were placed respectively on the forehead and tail of the animals. During the recordings, mice were lying down on a homeothermic blanket (Harvard Apparatus, Holliston, MA) to maintain body temperature around 37°C.

Typical ERG waveforms exhibiting a- and b-waves amplitude and implicit time in photopic and scotopic conditions are presented in **Supplemental Figure 1**. For every animal, rods’ function (scotopic) was first measured followed by cones’ function (photopic) using an Espion E1 system with flash stimulation provided by a Ganzfeld ColorDome (Diagnosys LLC, Lowell, MA). ERG signals were recorded using the Espion 3.0.1 software version (Diagnosys LLC, Lowell, MA). Prior to the scotopic ERG, mice’ eyes were adapted to dark for 1 hour. A light adaptation period of 10 minutes at 30 cd/m^2^ was allowed before photopic recordings. Rods’ function was assessed using seven white flashes of luminance increasing from -0.020 to 2.859 log cd.s/m^2^ produced by the Color Dome LEDs. Interstimulus intervals were 15 s for the first intensity and 30 s for the last 6 highest intensities. Color Dome xenon flashes were used for photopic ERG, using five increasing flashes ranging from 0.885 to 2.859 log cd.s/m^2^ with interstimulus intervals set at 30 s. For both scotopic and photopic ERG, at least four responses were recorded at each intensity.

**Figure 1.**
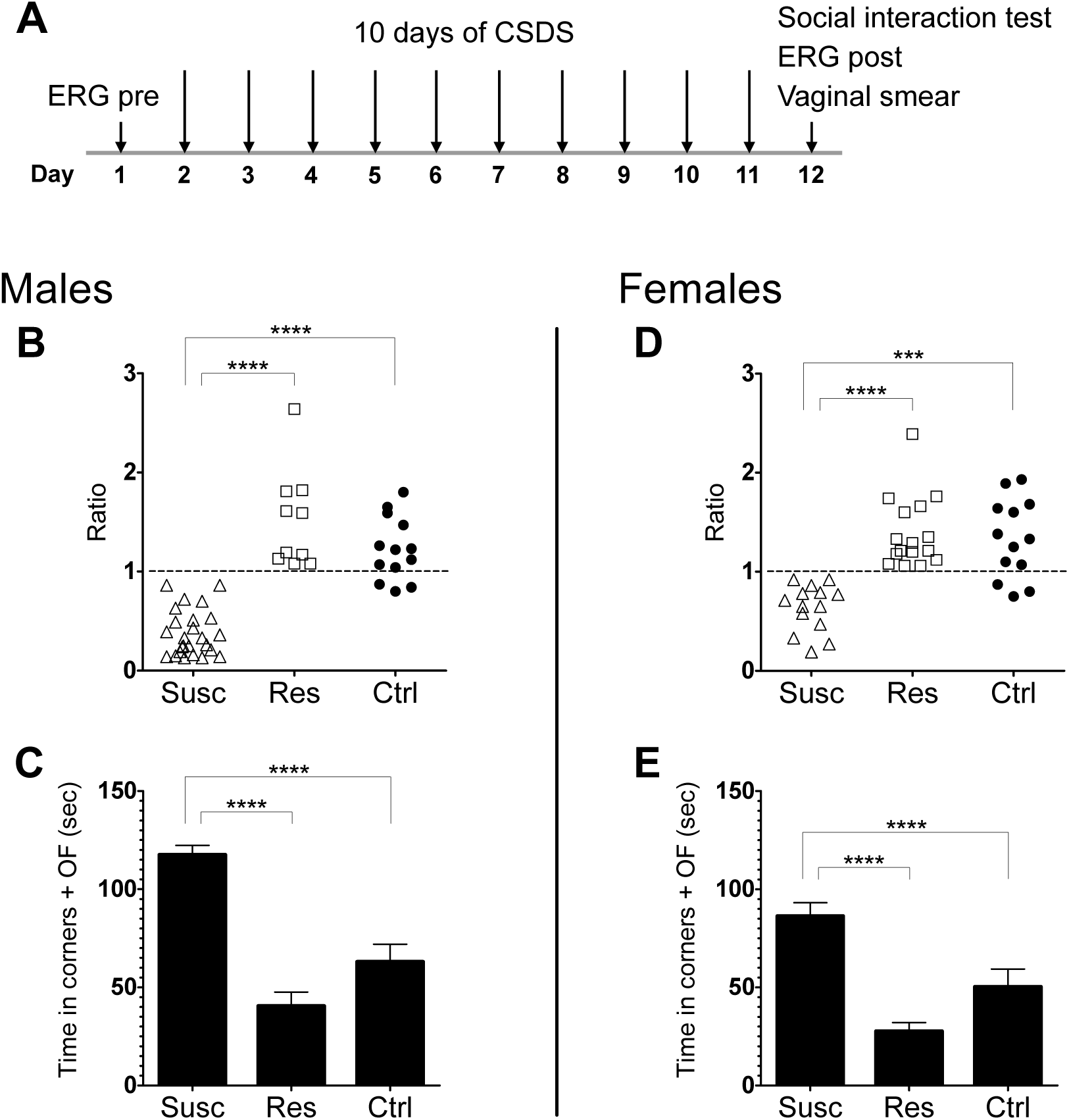
**(A)** Schematic representation of the experimental timeline. Pre-stress ERG was assessed at day 1, followed by 10 days of chronic social defeat stress (CSDS) 24 hours after (day 2, ending day 11). The social interaction test and the post-stress ERG were assessed 24 hours after the last defeat at day 12. (**B, D**) Susceptible, resilient, and control mice are divided according to their social interaction (SI) ratio score (time in interaction zone with social target CD-1 / time in interaction zone without the social target). Male (**B**) and female (**D**) mice susceptible to CSDS exhibit social avoidance compared to male and female resilient and control mice. (**C, E**) Male (**C**) and female (**E**) susceptible mice spent significantly more time in the corner and open field zones than the male and female resilient and control mice. Significance was determined using two-way ANOVA with Bonferroni multiple comparisons test. Symbols and bars represent the group average ± SEM; * *p* < .05, ** *p* < .01, *** *p* < .001; *n*-values are given in the method section.

Baseline ERG measures were collected for every mouse one day before the beginning of defeat stress and repeated 24 hours after the last defeat (**figure 1A**). Vaginal smears were collected before each ERG recording to determine estrous cycles for every female mouse (**Supplemental Table 1**).

### Data Analysis

To test whether, in both males and females, susceptible mice interacted less with the target CD-1 than the resilient and control mice, a two-way ANOVA with a Bonferroni correction was used to compare the social interaction (SI) ratios of each sex and phenotype. As for the difference between the three phenotypes in avoiding the target CD-1, time spent in the corners and the interaction zone during the second phase of the SI test was compared using the same analysis model. In order to investigate if the estrous cycle has repercussion on the phenotype of the female mice, a one-way ANOVA (Kruskal-Wallis test for non-parametric data) was used to compare the SI ratios of the females according to their estrous phase (estrus, metestrus, diestrus, pro-estrus) measured the same day as the social interaction test.

For both scotopic and photopic recordings, mean a- and b-waves amplitudes and implicit times for every step were calculated using Espion 3.0.1. Outliers for each group were then identified using the GraphPad’s QuickCalcs Grubb’s test (GraphPad Software Inc., San Diego, USA) and removed when applicable. All statistical analyses were performed on the Vmax (the intensity of light at which the retina saturates) of each ERG parameters with SPSS Statistics 26.0 (IBM Corp., Armonk, USA). Scotopic and photopic ERGs were analyzed separately. Scotopic and photopic Vmax values were set at 1.89 and 2.39 log cd.s/m^2^ according to their respective luminance response function (LRF) (**Supplemental figure 2**).

**Figure 2.**
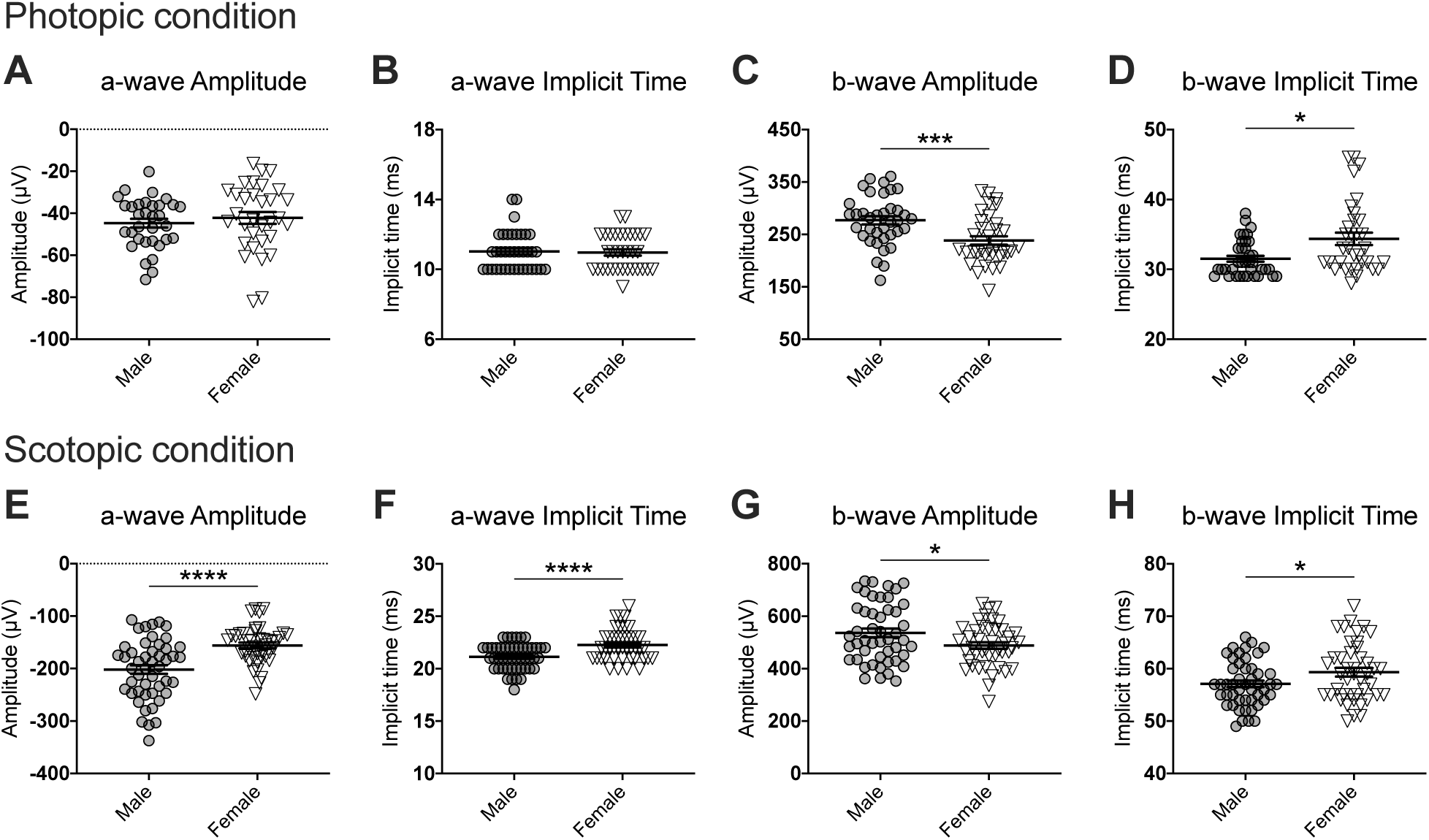
Comparisons of the male and female ERG parameters at baseline for photopic **(A-D)** and scotopic **(E-H)** conditions. Overall, males at baseline exhibit stronger and faster signals than females, excepted for the photopic a-wave amplitude **(A)** and implicit (me **(B)**. These results establish that retinal activity at baseline is different in males and females. Significance was determined using two-way ANOVA with LSD multiple comparisons test. Scatter plots depict each sample value of all groups of mice and bars represent the group average ± SEM; * *p* < .05, ** *p* < .01, *** *p* < .001, **** *p* < .0001; *n*-values are given in the method section.

ERG data are presented as the mean and standard error of each parameter. After checking the normality assumption using Shapiro-Wilk test, a two-way ANOVA was used to compare male and female susceptible, resilient, and control mice at baseline. Variations in ERG parameters for each sex were then tested separately using two-way mixed ANOVA models in which phenotype (susceptible, resilient and control) and time of assessment (pre-stress versus post-stress) were included as main factors. Our model examined the two-way interactions between phenotype and time. Post hoc were analyzed using Fisher’s least significant difference (LSD) multiple comparisons test. All P-values under .05 were considered as significant.

Multiple backward logistic regressions were used to obtain the best combination of ERG parameters and sex as covariates to predict post-stress phenotype in males and females. The accuracy of the model was calculated with the area under the receiver operating characteristic (AU-ROC) curves: an AU-ROC curve of 1 corresponded to a model with perfect discrimination ability while an AU-ROC of 0.5 corresponded to a model with no discrimination ability (meaning that the probability of having the predicted phenotype would equal chance). Sensitivity and specificity were estimated by comparing the predicted group of each mouse to their true group.

## Results

In this study, we performed scotopic and photopic ERGs before (pre) and after (post) chronic social defeat stress (CSDS) in male and female mice (**figure 1A**). The control mice ERGs were also assessed during the pre- and post-stress without being submitted to CSDS. Under this experimental design, we tested with the ERG (1) whether susceptible and resilient mice exhibited specific changes in the activity of retinal cells; (2) whether social stress changes retinal cells activity differently in males and females; and (3) whether measures before stress (pre) could predict which male and female mice would become susceptible or resilient to social defeat stress.

First, our results show that CSDS indeed induced the expression of stress susceptibility and resilience in both males and females. Precisely, 10 days of CSDS induced social avoidance in 27 males and 15 females while the remaining 8 males and 15 females continued to interact with CD1 targets (significant effect of phenotypes: *F*_(2, 87)_ = 56.7, *p* < .0001; **figure 1 B, D**). As expected, susceptible male and female mice rather spent significantly more time in the corners and the open field avoiding CD1 targets compared to control (*p* < .0001) and resilient mice (*p* < .0001) (**figure 1 C, E**). Noticeably, our analysis detected no effect of the estrous cycle on SI ratio in either susceptible, resilient, or control mice (χ^2^ = 1.38, *p* = .711, df = 3; **Supplemental Table 1**).

### Sex-specific retinal activity measured with ERG

One of the main objectives of this study was to test whether photopic and scotopic retinal activities vary in males and females. To answer this question, we first used a two-way ANOVA with LSD multiple comparisons test to compare male and female ERGs at baseline. Overall, for most ERG parameters, our analyses revealed a significant main effect of sex, with post hoc tests suggesting that males have elevated amplitudes and faster implicit times than females in both photopic (a-wave amplitude: *F*_(1, 59)_ = 0.62, *p* > .05; a-wave implicit time: *F*_(1, 65)_ = 0.01, *p* > .05; b-wave amplitude: *F*_(1, 65)_ = 14.1, *p* < .001; b-wave implicit time: *F*_(1, 65)_ = 4.49, *p* < .05; **figure 2 A-D**) and scotopic (a-wave amplitude: *F*_(1, 85)_ = 21.7, *p* < .0001; a-wave implicit time: *F*_(1, 85)_ = 19.6, *p* < .0001; b-wave amplitude: *F*_(1, 85)_ = 6.37, *p* < .05; b-wave implicit time: *F*_(1, 85)_ = 6.82, *p* < .05; **figure 2 E-H**) conditions. Because of this important sexual dimorphism, male and female ERG activity following social defeat was analyzed separately.

### Variation in retinal activity in mice becoming susceptible or resilient to social defeat stress

We next tested the impact of CSDS on photopic and scotopic signals by using a two-way mixed model ANOVA with timing (pre-vs post-stress) and mouse phenotype (control, susceptible and resilient) as main factors (See **Supplemental Table 2-3** for a summary of all statistical outputs). For both males and females, our analyses of the photopic condition revealed no significant effect of time by phenotype interaction for all of the parameters (all *p* > .05) (**Supplemental Table 2; Supplemental figure 3-4**) suggesting that 10 days of social defeat has no functional impact on the cone activity in males and females.

In contrast, our analysis of the scotopic condition revealed significant time by phenotype interactions in males and females. In male mice, analysis of the a-wave amplitude revealed a significant time by phenotype interaction (*F*_(2, 75.41)_ = 3.88, *p* < .05; **Figure 3 A-C**) with post hoc analyses highlighting a significant decrease of amplitude in susceptible (−16.6%; *p* < .05) and resilient (−37.2%; *p* < .001) mice after the CSDS. This decrease resulted in a significant smaller amplitude in susceptible (*M* = -165.54, *SE* = 6.78; *p* < .05) and resilient (*M* = -142.30, *SE* = 13.07; *p* < .01) mice compared to control (*M* = -192.87, *SE* = 9.59) mice after CSDS. Similarly, analysis of the a-wave implicit time (time by phenotype interaction: *F*_(2, 84.32)_ = 4.67, *p* < .05; **Figure 3 D-F**) suggested that 10 days of social defeat shortens the implicit time in susceptible mice (−0.81 ms; *p* < .01), but not in resilient mice (−0.09 ms; *p* > .05), whereas control mice showed a non-significant pre-post change of - 0.64 ms (*p* > .05). Interestingly, the susceptible (*M* = 21.58, *SE* = 0.23) mice had a significant prolonged a-wave implicit time at baseline when compared to the resilient (*M* = 20.63, *SE* = 0.41; *p* < .05) and control (*M* = 20.57, *SE* = 0.31; *p* < .05) mice. Finally, analysis of the b-wave amplitude identified a significant main effect of time (*F*_(1, 68.99)_ = 5.02, *p* < .05; **Figure 3 G-I**), supporting a general pre-post decrease of 9.52% in all groups, but no significant phenotype or time by phenotype interaction main effect. A similar lack of effect was revealed for b-wave implicit time (**Figure 3 J-L)**. To summarize the main findings in male, the susceptible mice showed a prolonged a-wave implicit time at baseline and demonstrate a shorten a-wave implicit time after CSDS. The a-wave is depressed in both susceptible and resilient phenotypes after CSDS while the b-wave is depressed in all groups during the post-stress assessment.

**Figure 3.**
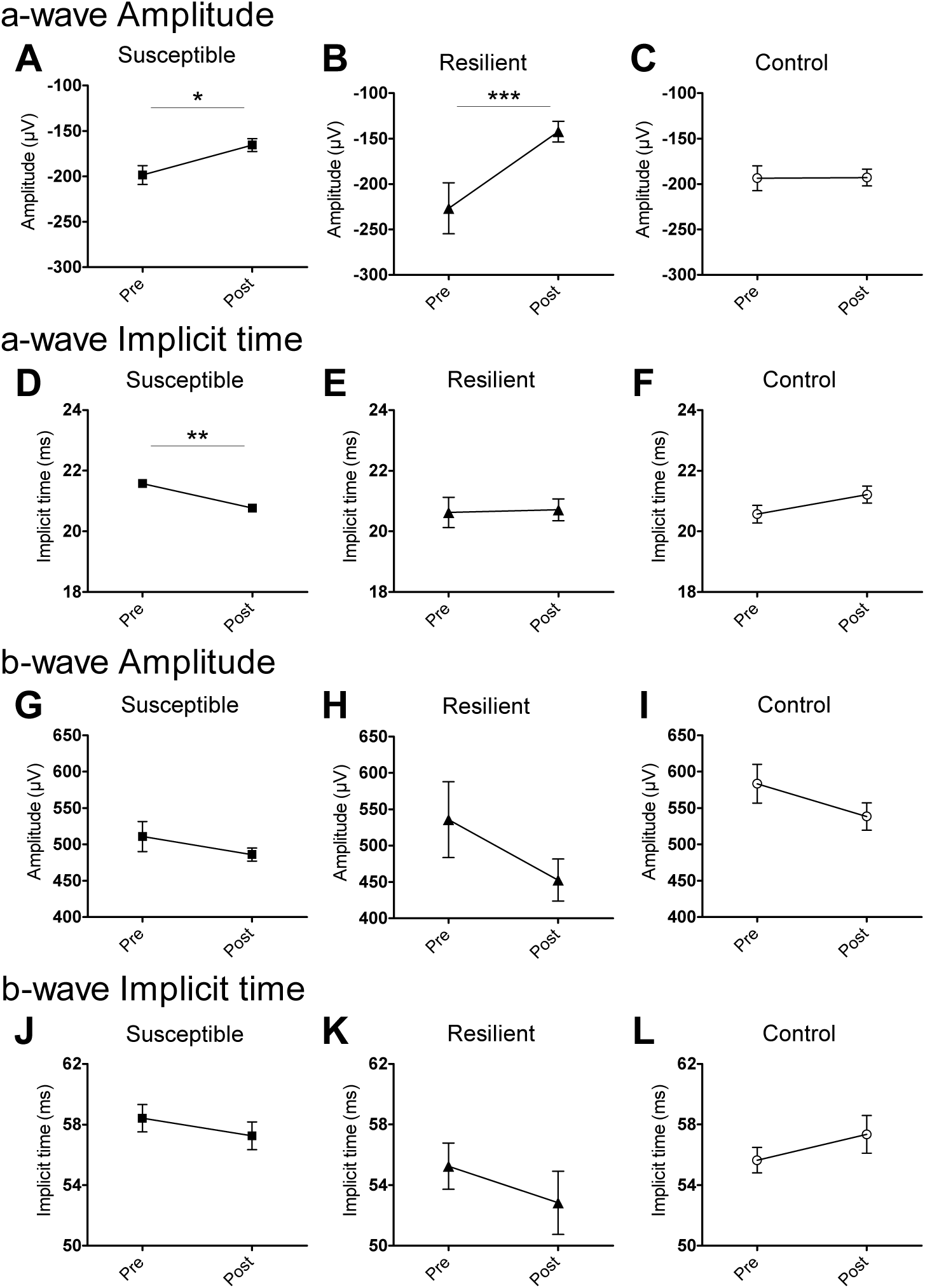
The evolution of each ERG parameters under the scotopic condition at baseline and after CSDS in male mice. The susceptible, resilient, and control male a-wave amplitude **(A-C)**, a-wave implicit time **(D-F)**, b-wave amplitude **(G-I)**, and b-wave implicit time **(J-L). (A-C)** The time by phenotype interaction in the a-wave amplitude revealed a sex-specific signature of the stressed males. **(D-F)** The time by phenotype interaction in the a-wave implicit time is associated with susceptibility to stress in males. Significance was determined using two-way mixed ANOVA with LSD multiple comparisons test. Symbols and bars represent the group average ± SEM; * *p* < .05, ** *p* < .01, *** *p* < .001; *n*-values are given in the method section.

In females, our analysis of the a-wave amplitude in scotopic condition revealed a significant main effect of time (*F*_(1, 75.97)_ = 16.4, *p* < .001; **Figure 4 A-C**) showing that a-wave amplitude is depressed in susceptible, resilient and control mice (pre vs post values for each group). Interestingly, our analysis highlighted a trend toward a significant time by phenotype main effect for the a-wave implicit time (*F*_(2, 67.78)_ = 2.68, *p* = .076; **Figure 4 D-F**). Subsequent post hoc tests suggest that the post a-wave implicit time is significantly prolonged in control mice (+1.41 ms; *p* < .05) when compared to baseline, but not in those becoming susceptible (−0.75 ms; *p* > .05) or resilient (−0.10 ms, *p* > .05). Furthermore, the post a-wave implicit time is significantly prolonged in control mice (*M* = 23.42, *SE* = 0.56) when compared to susceptible mice (*M* = 21.64, *SE* = 0.52; *p* < .05). Finally, we identified a significant main effect of time (*F*_(1, 76.48)_ = 7.79, *p* < .01; values pre versus post for every group; **Figure 4 G-I**) in the b-wave amplitude showing an overall decrease of 10.6% in all groups, and found no significant main effect in the b-wave implicit time (**Figure 4 J-L**). In summary, similarly to the susceptible male mice, female mice tend to demonstrate a shorter a-wave implicit time after CSDS, but the difference does not reach significance. Combined with the delay observe in control mice, this leads to a significant post a-wave implicit time difference between control mice and susceptible mice.

**Figure 4.**
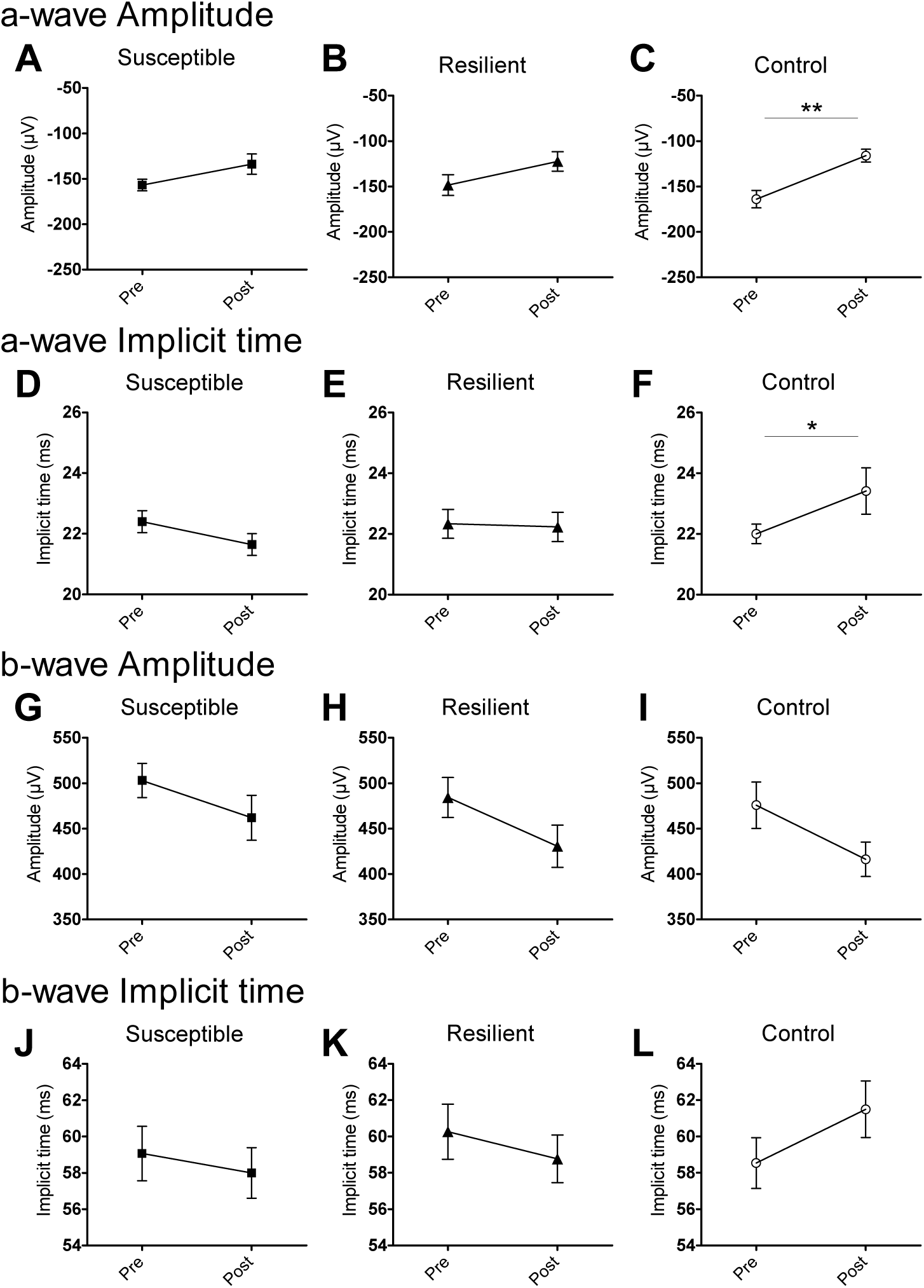
The evolution of each ERG parameters in the scotopic condition at baseline and after CSDS in female mice. The susceptible, resilient, and control female a-wave amplitude **(A-C)**, a-wave implicit time **(D-F)**, b-wave amplitude **(G-I)**, and b-wave implicit time **(J-L). (D-F)** The time by phenotype interaction in the a-wave implicit time is associated with susceptibility to stress in females. Significance was determined using two-way mixed ANOVA with LSD multiple comparisons test. Symbols and bars represent the group average ± SEM; * *p* < .05, ** *p* < .01, *** *p* < .001; *n*-values are given in the method section.

### Predictive ability of the ERG

Finally, we tested our capacity to predict the expression of stress susceptibility and resilience based on baseline ERG signals using receiver operating characteristic (ROC) curves combined with multiple backward logistic regressions. From these analyses, our results suggest that none of the fitted models reached significance. However, the backward procedure highlighted a trend toward significance in the model fitting the a-wave implicit time using sex as a covariate (*p* = .0537). The AU-ROC curve of this model reaches 0.6941 (**Figure 5A**) – which is considered as a weak, but acceptable predictive model – with a sensitivity and specificity of 80.5 and 39.1, respectively (**Figure 5B**). Even if the model combining all four ERG parameters with sex has an estimated AU-ROC curve of 0.7084, it is not significantly different than 0.05 (*p* > .05). The same applies for the estimated AU-ROC curves of the a- and b-wave amplitudes (0.6479; and 0.6278 respectively) and the b-wave implicit times (0.6718) separately when combined with sex (all *p* > .05) (F**igure 5B**). Overall, results from this analysis suggest that the ERG has a weak capacity to predict the expression of susceptibility or resilience phenotypes in males or females subjected to social defeat.

**Figure 5.**
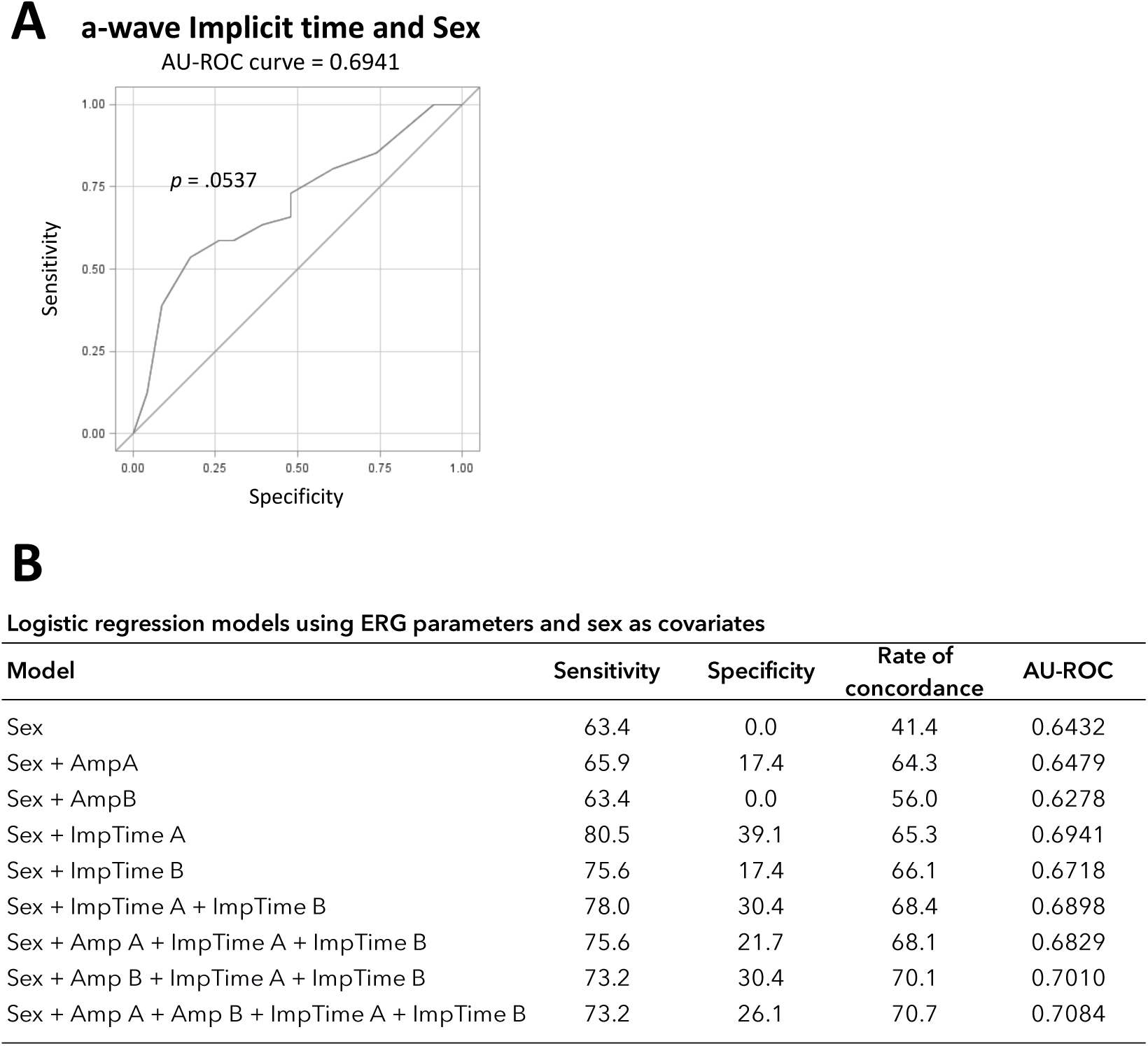
Capacity of the ERG in predicting the expression of stress susceptibility and resilience from the baseline signals.**(A)** The fitted model of the a-wave implicit time with sex is near significantly different than 0.05 (*p* = .0537) with an AU-ROC curve of 0.6941. **(B)** Table presenting the different fitted models analysed by the multiple backward logistic regressions with their sensitivity, specificity, rate of concordance, and area under the receiver operating characteristic (AU-ROC) curves. Statistical significance is set at *p* < .05; *n*-values are given in the method section.

## Discussion

In this study, we tested if the phenotypes of susceptibility and resilience could be associated with specific ERG alterations and whether baseline ERG signals could detect which males and females undergoing CSDS would develop such phenotypes. Our results show that CSDS induces changes in the activity of the retina that reached significance in males but not in females. Additionally, our analysis suggests that baseline ERG measures, based on our relatively small sample size, might not be sufficient to predict mice’s phenotype to social stress.

Our findings suggest that stress susceptibility in males associates with shorter a-wave implicit times following CSDS. Since the a-wave represents the activity of the photoreceptors^44^, this alteration might correspond to a faster hyperpolarization of the rods following light stimulus. This variation of the a-wave implicit time might also be the result of a faster intrusion of the b-wave in the ERG, thus reflecting a faster ON-bipolar cells depolarization following the decrease in glutamate release from the rods^45^. This alteration indeed testifies not only as a specific manifestation of social stress, but also of susceptibility to stress in males. Similarly, our results suggest that social stress induces a depressed a-wave amplitude in susceptible and resilient male mice, likely implying a reduction in the hyperpolarization of the rods in response to light stimulus. In contrast, results regarding stress susceptibility in females were not conclusive enough to associate with specific ERG features to CSDS. This difference might be explained by the fact that the number of susceptible mice in our analysis was almost twice larger in males than females. Besides, our main findings in females relate to the presence of an aging effect, which could have interfered with the final outcomes. Regarding the b-wave implicit time and amplitude, no effect of social stress was detected in males and females in both susceptible and resilient mice suggesting that the activity of the ON-bipolar cells is maintained after social stress. The lack of specific ERG alteration in resilient mice is surprising given that resilience to social stress is perceived as an adaptative mechanism involving molecular activations in pathways allowing mice to cope with chronic stress and exhibit adaptative behavioral responses^43,46^. Overall, our findings are consistent with the notion that social stress induces a wide range of adaptative responses^47,48^, not only in the brain, but also in peripheral organs and support the use of ERG to detect these alterations in the retina of males susceptible to social stress.

Monoamines like dopamine (DA) or serotonin (5-HT) are known to act as a central player in the neurochemical imbalance hypothesis underlying the pathophysiology of depression and stress-induced behavioral deficits^49,50^. Interestingly, previous studies using DAT-KO and Tph2-KI mice showed that alterations in central monoamines impact the ERG^37^. Likewise, ERG alterations found in MDD patients tend to normalize following SNRIs antidepressant treatments^30^ modulating central DA, 5-HT, and norepinephrine neurotransmission^51^. Together, these observations suggest that changes in central monoamines signaling can modulate the activity of the retina which is consistent with our findings in mice susceptible to CSDS. However, unlike our results, most of the behavioral deficits found in these studies were associated with photopic alterations. This difference might be explained by the use of a stress paradigm instead of transgenic mice or pharmacotherapy which might lead to important molecular and physiological adaptations. This being said, the mechanisms linking alterations of central DA and 5-HT circuits with the activity of the retina are still unclear. However, evidences of direct histaminergic and serotonergic retina projections from the brain (also referred as retinopetal, centrifugal, or efferent neurons) have been reported in mammals^52-54^. In mice, retinal dopaminergic cells – mostly the amacrine cells^55^ – are among the targets of the histaminergic retinopetal axons^56,57^. Since light adaptation in the retina is modulated by DA^58^ and that changes in central DA can affect histamine release^59,60^, the ERG alterations associated with stress susceptibility in our study might be accounted, in part, by the influence of the histaminergic retinopetal axons. 5-HT also acts as a neuromodulator in the retina^61^ and its activity could potentially be modulated by serotonergic retinopetal neurons emerging from the dorsal raphe nuclei (DRN)^53,62,63^. Not only it has been reported that bilateral administration of the neurotoxic agent 5-7-dihydroxytryptamine into the DRN decreases 5-HT and SERT content in the retina^62^, electrophysiological stimulation of the DRN in rats have been shown to induce specific ERG alterations^63^. It is therefore legitimate to assume a relationship between the serotonergic retinopetal projections and the ERG alterations found in susceptibility to stress. Finally, given that CSDS was performed over 10 days, ERG alterations found in our study may be the result of a slow-paced process. Accordingly, since photoreceptors employ various protein isoforms and require several components and neuromodulators for the integrity of their functions, one could argue that retinal activity may be modulated by its connections with the retinal pigment epithelium (RPE)^64,65^ which could be promoted in susceptible mice by the changes induced by social stress on the permeability of the blood brain barrier^66,67^, or in this case, the blood retina barrier.

Our findings also allowed us to reveal the existence of a sexual dimorphism in the activity of the retina under basal condition and following social stress. Our findings suggest that the baseline ERG amplitudes are roughly 15% bigger and 4% faster in males than in females under both scotopic and photopic conditions suggesting that males and females may be processing retinal information differently. However, these results are divergent than those observed by Mazzoni, et al. ^68^ who reported no differences between male and female C57BL6 wild-type mice at the same age. Regardless, considering that many sex-specific features were identified in gene expression throughout the lifespan of the mouse^69^ and that plasticity of the visual system still occurs at this period of age in mice^70,71^, it should not be inconceivable to find significant differences between male and female basal ERG responses. Another possibility explaining male and female ERG difference relates to variations in hormonal status^72^ although no differences in female estrous cycles were found in our study, reducing the impact of this potential factor. Regarding stress susceptibility, our findings suggest that males susceptible to social stress exhibit explicit retinal alterations following stress whereas it is not as conclusive in females. This lack of effect in females specifically is surprising as a growing body of evidence describes distinct physiological^76,77^, functional^78^ and molecular^79^ alterations induced by chronic stress in males and females. It is also believed that these differences may be part of the reasons underlying why males and females respond differently to stress^11,12^. The lack of effect in females after CSDS observed in our study, however, is likely due to the small sample size in females. Future work using the ERG in the context of stress should consider using larger sample size in order to detect reliable signal in females more specifically.

Past and recent clinical studies highlighted the fact that no reliable biomarkers or specific endophenotypes exist for MDD and associated diseases^18,21,23,49,50^. A similar lack of predictive capabilities exists in preclinical research using animal models. Here, we provided results suggesting that the ERG might predict mouse phenotype before the social stress with the efficiency of 69.5% using the a-wave and sex as predictors. Even though this efficiency is considered as weak and that this fitted model is not yet significant (*p* = .0537), this remains a promising outcome. This is indeed in line with previous results using ERG from which SZ patients were differentiated from individuals suffering from BP disorder with 86% accuracy^32^. Importantly, our results, along with those in clinical research using ERG^29,30,33,35^, suggest that retinal activity might represent an endophenotype which could be used to determine whether mice or human facing social stress are more likely to develop specific aspects of the disease. This is of the upmost importance as by combining the predictive capabilities of ERG with molecular or functional studies, one could specifically target some of the systems directly involved in the development of stress resilience or susceptibility while the animals are developing these phenotypes. This being said, our statistical model was limited by our sample size which did not allow us to reach sufficient power to establish robust predictions.

It should also be noticed that, in contrast with diurnal humans, mice are nocturnal mammals with different spectral sensitivity inherent to the distinct population and density of photoreceptors in their eyes^80^. These distinctions might explain some of the differences observed in our studies and previous ERG analyses performed in clinical human population^30,33^. This consideration may become even more important considering that most significant results in humans originate from light-adapted (photopic) ERGs^30,33^ while most of our results were obtained in the dark-adapted condition (scotopic). These facts should be considered when interpreting the translational value of our findings in a mouse model of chronic social stress.

Overall, the search for reliable biomarkers and endophenotypes of stress responses has been shaping clinical and preclinical research for the last decades with still limited success^18,21,23,49,50^. Here, we present results suggesting that not only social stress induces changes in retinal activity that associate with the expression of susceptibility in a mouse model of CSDS but also that ERG can be used to predict the mouse phenotype to social stress. This being said, CSDS is one of many mouse models^81,82^ and previous molecular studies highlighted the capacity of different mouse models to reproduce not only the clinical manifestations but also the molecular changes relevant to MDD in males and females^79,82^. Thus, it will be important to determine whether different types of stress induce similar retinal changes and whether ERG can be used to predict the expression of different behavioral alterations triggered by the distinct behavioral paradigms used to model depression in males and females. Nevertheless, the exciting options offered by the ERG in preclinical research pave the way to novel research avenues for studying the molecular and functional mechanisms underlying the expression of stress responses and develop new therapeutic options to treat MDD more efficiently.

## Acknowledgements

B.L. holds a Sentinelle Nord Research Chair, is supported by a NARSAD young investigator award, a CIHR (SVB397205), and Natural Science and Engineering Research Council (NSERC; RGPIN-2019-06496) grants and receives FRQS Junior-1 salary support; this work was also made possible by resources supported by the Quebec Network on Suicide, Mood Disorders and Related Disorders. M.H. holds a CIHR grant (MOP-82707).

## Author contributions

B.L., M.H., and E.A. conceived and designed this research. E.A. and A.A.L. performed the experiments and ran the behavioral testing. E.A., S.M., and K.F. carried out the analyses and prepared figures. A.A.L., T.B., F.Q., K.A, A.B., and E.A. performed tissue dissections. E.A. and B.L. wrote the manuscript.

