## Supplemental material for "Sex-specific retinal anomalies induced by chronic social stress in mice"

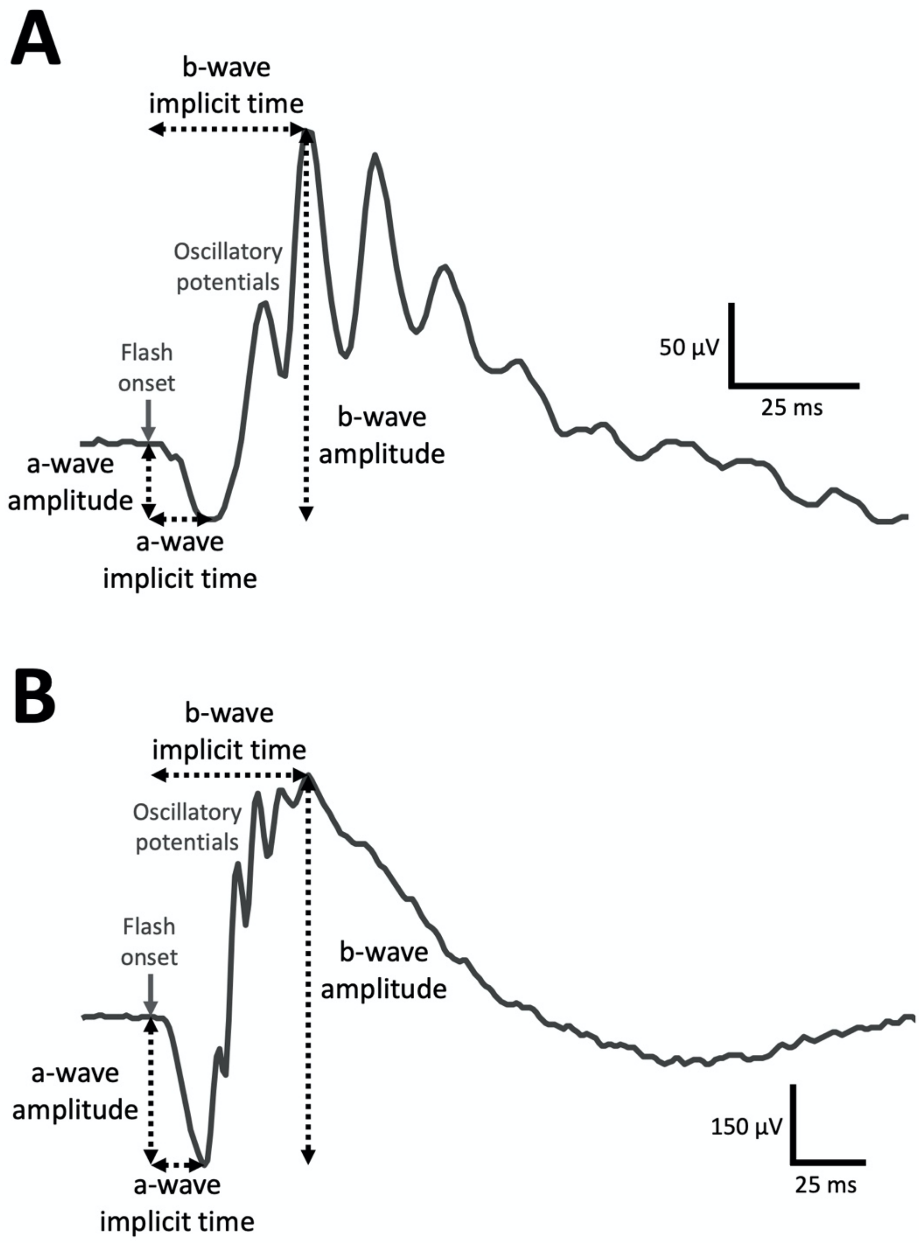


**Figure S1**
Example of a mouse ERG waveform obtained at Vmax under **(A)** photopic and **(B)** scotopic conditions. A- and b-wave amplitudes are calculated (in microvolts) from baseline to the bottom of the a-wave and, from there, to the peak of the b-wave, respectively. Implicit time corresponds to the total time (in milliseconds) from the flash onset until the bottom of the a-wave or the peak of the b-wave are reached, respectively.

**Supplemental table 1 | Distribution of the social interaction ratios upon the estrous cycle of the susceptible, resilient, and control mice**

| | Susceptible | | Resilient | | Control | | Total | |
|---|---|---|---|---|---|---|---|---|
| | *n* | *M* | *n* | *M* | *n* | *M* | *n* | *M* |
| Estrus | 5 | .579 | 6 | 1.33 | 7 | 1.07 | 18 | 1.02 |
| Metestrus | 5 | .720 | 4 | 1.32 | 4 | 1.41 | 13 | 1.12 |
| Diestrus | 3 | .699 | 0 | - | 1 | 1.89 | 4 | .998 |
| Pro-estrus | 3 | .586 | 4 | 1.64 | 1 | 1.64 | 8 | 1.35 |

**Table S1**
Distribution of the SI ratios upon the estrous cycle of the susceptible, resilient, and control female mice. The SI ratio is calculated from the time spent in the interaction zone (time in interaction zone with social target CD-1 / time in interaction zone without the social target). *M* = mean.

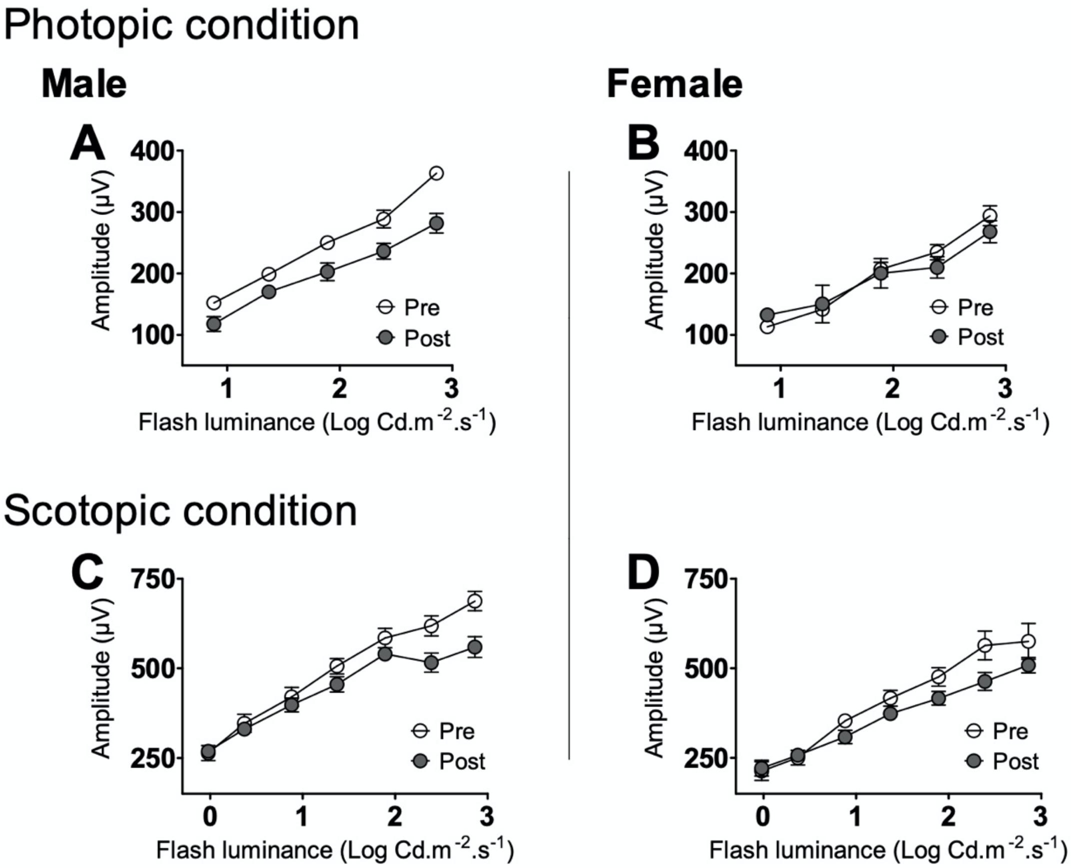


**Figure S2**
The b-wave data from scotopic and photopic conditions in males and females were plotted against the flash luminance used to generate the luminance response function (LRF) from which the Vmax is derived. Vmax refers to the maximal b-wave amplitude at which the activity of the retina saturates and corresponds to the first plateau in the LRF curves. Photopic and scotopic Vmax were set at 2.39 and 1.89 log of luminance, respectively. Symbols and bars represent the group average ± SEM.

**Supplemental table 2 | Two-way mixed ANOVA results of the photopic condition**

| Sex | Wave | Parameter | Time | Susceptible (M = 27, F = 15) | Resilient (M = 8, F = 15) | Control (M = 14, F = 13) | Fixed effet | | |
|---|---|---|---|---|---|---|---|---|---|
| | | | | *M (SE)* | *M (SE)* | *M (SE)* | *Phenotype P (F)* | *Time P (F)* | *Phenotype*Time P (F)* |
| **Male (*n* = 49)** | a-wave | Amplitude | Pre | -42.71 (3.01) | -43.77 (4.55) | -48.57 (3.80) | *.4597 (0.79)* | *.4408 (0.60)* | *.0969 (2.42)* |
| | | | Post | -41.13 (3.11) | -50.16 (5.39) | -36.05 (3.98) | | | |
| | | Implicit time | Pre | 10.95 (0.26) | 11.43 (0.43) | 10.91 (0.34) | *.9046 (0.10)* | *.2940 (1.12)* | *.2176 (1.56)* |
| | | | Post | 10.89 (0.25) | 10.33 (0.45) | 11.15 (0.31) | | | |
| | b-wave | Amplitude | Pre | 263.67 (10.63) | 294.66 (17.51) | 288.55 (13.97) | *.1353 (2.06)* | *< .0001 (21.2)* | *.8174 (0.20)* |
| | | | Post | 219.78 (10.22) | 233.02 (17.71) | 236.28 (12.03) | | | |
| | | Implicit time | Pre | 31.90 (0.77) | 33.43 (1.30) | 30.91 (1.04) | *.8808 (0.13)* | *.9380 (0.01)* | *.0622 (2.93)* |
| | | | Post | 31.83 (0.40) | 31.17 (0.69) | 33.08 (0.47) | | | |
| **Female (*n* = 43)** | a-wave | Amplitude | Pre | -39.13 (4.89) | -47.59 (4.89) | -39.71 (5.13) | *.2455 (1.45)* | *.9115 (0.01)* | *.9282 (0.07)* |
| | | | Post | -41.34 (5.74) | -49.25 (7.20) | -37.49 (7.77) | | | |
| | | Implicit time | Pre | 10.92 (0.38) | 10.82 (0.40) | 11.64 (0.40) | *.0605 (2.97)* | *.0792 (3.21)* | *.5542 (0.60)* |
| | | | Post | 11.91 (0.32) | 11.00 (0.38) | 12.20 (0.48) | | | |
| | b-wave | Amplitude | Pre | 230.01 (13.73) | 247.38 (14.34) | 238.56 (14.34) | *.5158 (0.67)* | *.2077 (1.63)* | *.4165 (0.89)* |
| | | | Post | 234.68 (9.87) | 231.51 (11.04) | 209.78 (12.75) | | | |
| | | Implicit time | Pre | 32.36 (1.50) | 36.25 (1.44) | 34.30 (1.57) | *.1753 (1.81)* | *.7049 (0.15)* | *.0599 (2.99)* |
| | | | Post | 33.22 (0.89) | 32.29 (1.01) | 36.20 (1.19) | | | |

**Table S2**

Summary of the two-way mixed model ANOVA results for the photopic condition comparing means and standard errors from susceptible, resilient, and control mice. The table also displays the fixed effect of each interaction from the analysis. See supplemental figures S3 and S4 for more details. *M* = mean; *SE* = standard error, *P* = p-value, *F* = Fisher's statistic. Significance is set at .05.

**Supplemental table 3 | Two-way mixed ANOVA results of the scotopic condition**

| Sex | Wave | Parameter | Time | Susceptible (M = 27, F = 15) | Resilient (M = 8, F = 15) | Control (M = 14, F = 13) | Fixed effet | | |
|---|---|---|---|---|---|---|---|---|---|
| | | | | *M (SE)* | *M (SE)* | *M (SE)* | *Phenotype P (F)* | *Time P (F)* | *Phenotype*Time P (F)* |
| **Male (*n* = 49)** | a-wave | Amplitude | Pre | -198.55 (11.21) | -226.76 (20.21) | -193.64 (15.28) | *.6017 (0.51)* | *.0006 (13.0)* | *.0249 (3.88)* |
| | | | Post | -165.53 (6.78) | -142.30 (13.07) | -192.87 (9.59) | | | |
| | | Implicit time | Pre | 21.58 (0.23) | 20.63 (0.41) | 20.57 (0.31) | *.2001 (1.64)* | *.9168 (0.01)* | *.0119 (4.67)* |
| | | | Post | 20.77 (0.17) | 20.71 (0.33) | 21.21 (0.24) | | | |
| | b-wave | Amplitude | Pre | 510.88 (21.86) | 535.94 (39.41) | 583.51 (29.79) | *.0092 (5.02)* | *.0151 (6.21)* | *.5186 (0.66)* |
| | | | Post | 486.10 (11.27) | 452.76 (21.73) | 538.48 (15.94) | | | |
| | | Implicit time | Pre | 58.42 (0.82) | 55.25 (1.48) | 55.64 (1.12) | *.0216 (4.01)* | *.5639 (0.34)* | *.2767 (1.30)* |
| | | | Post | 57.26 (0.92) | 52.83 (1.94) | 57.36 (1.27) | | | |
| **Female (*n* = 43)** | a-wave | Amplitude | Pre | -156.91 (9.16) | -148.37 (9.16) | -164.01 (9.84) | *.5781 (0.55)* | *.0001 (16.4)* | *.3993 (0.93)* |
| | | | Post | -133.89 (9.54) | -122.42 (10.24) | -116.06 (10.66) | | | |
| | | Implicit time | Pre | 22.40 (0.39) | 22.33 (0.39) | 22.00 (0.42) | *.3579 (1.04)* | *.6340 (0.23)* | *.0758 (2.68)* |
| | | | Post | 21.64 (0.52) | 22.23 (0.54) | 23.42 (0.56) | | | |
| | b-wave | Amplitude | Pre | 502.97 (21.54) | 484.34 (21.54) | 475.85 (23.14) | *.2560 (1.39)* | *.0066 (7.78)* | *.9133 (0.09)* |
| | | | Post | 461.98 (21.67) | 430.62 (23.27) | 416.18 (24.22) | | | |
| | | Implicit time | Pre | 59.07 (1.45) | 60.27 (1.45) | 58.54 (1.55) | *.5810 (0.55)* | *.9115 (0.01)* | *.2636 (1.36)* |
| | | | Post | 58.00 (1.34) | 58.77 (1.44) | 61.50 (1.50) | | | |

**Table S3**

Summary of the two-way mixed model ANOVA results for the scotopic condition comparing means and standard errors from susceptible, resilient, and control mice. The table also displays the fixed effect of each interaction from the analysis. See figures 3 and 4 for more details. *M* = mean; *SE* = standard error, *P* = p-value, *F* = Fisher's statistic. Significance is set at .05.

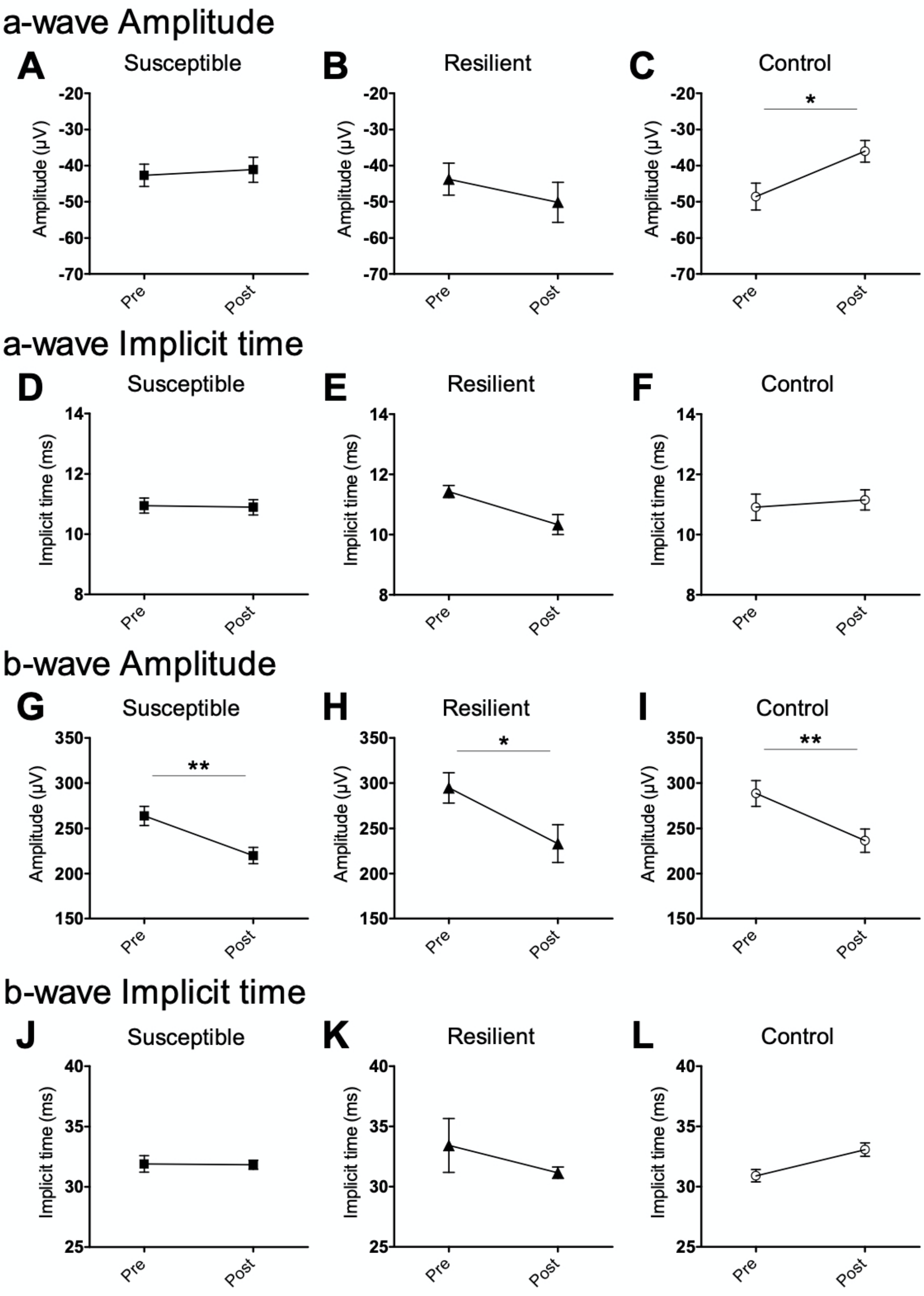
a-wave Amplitude
A
Susceptible
B
Resilient
C
Control
Amplitude (µV)
-20
-30
-40
-50
-60
-70
*
Pre
Post
a-wave Implicit time
D
Susceptible
E
Resilient
F
Control
Implicit time (ms)
14
12
10
8
Pre
Post
b-wave Amplitude
G
Susceptible
H
Resilient
I
Control
Amplitude (µV)
350
300
250
200
150
**
*
**
Pre
Post
b-wave Implicit time
J
Susceptible
K
Resilient
L
Control
Implicit time (ms)
40
35
30
25
Pre
Post

**Figure S3**
The evolution of each ERG parameters under the photopic condition at baseline and after CSDS in male mice. **(A-F)** The analyses in males reveal no significant interaction nor main effect in the a-wave amplitude (A-C) and implicit time (D-F). **(G-I)** No interaction found in the b-wave amplitude, however the analyses shows a main effect of time ($F_{(1, 67.87)} = 21.15$, $p < .0001$) revealing an average decrease of 52.6 µV at post-stress in all the groups of male mice (not showed). **(J-L)** The time by phenotype interaction is near significant ($F_{(2, 51.55)} = 2.932$, $p = .0622$) in the b-wave implicit time, however the post hoc depicts no further interaction. Significance was determined using two-way mixed model ANOVA with LSD multiple comparisons test. Symbols and bars represent the group average ± SEM; * $p < .05$, ** $p < .01$, *** $p < .001$; $n$-values are given in the method section.

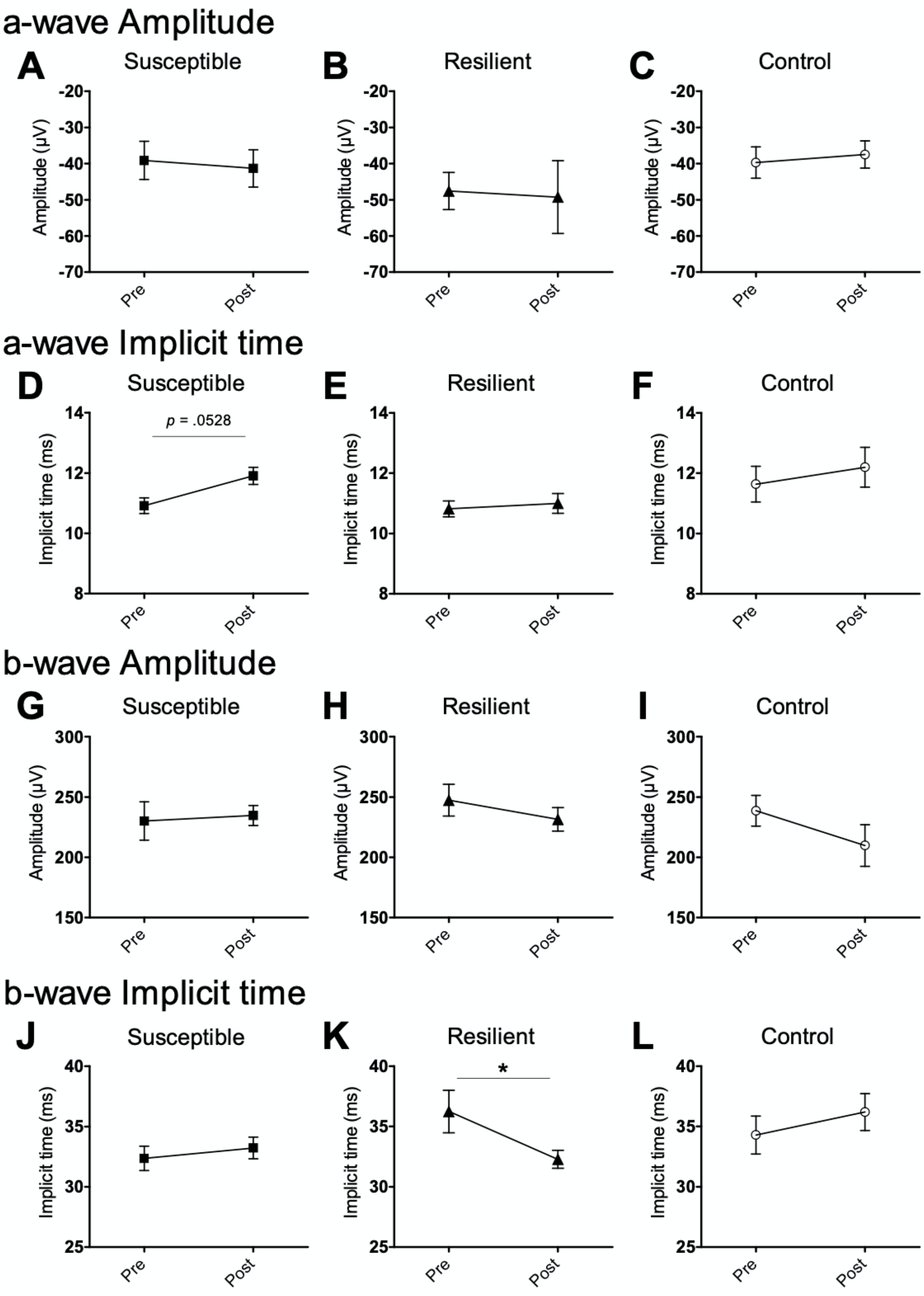
a-wave Amplitude
A
Susceptible
B
Resilient
C
Control
Amplitude (μV)
-20
-30
-40
-50
-60
-70
Pre
Post
a-wave Implicit time
D
Susceptible
p = .0528
E
Resilient
F
Control
Implicit time (ms)
14
12
10
8
b-wave Amplitude
G
Susceptible
H
Resilient
I
Control
300
250
200
150
b-wave Implicit time
J
Susceptible
K
Resilient
*
L
Control
40
35
30
25

**Figure S4**

The evolution of each ERG parameters in the photopic condition at baseline and after CSDS in female mice. **(A-I)** The analyses in females reveal no significant interaction nor main effect in the a-wave amplitude (A-C) and implicit time (D-F), and the b-wave amplitude (G-I). **(J-L)** Analysis of the b-wave implicit time however revealed a trend in the time by phenotype interaction ($F_{(2, 47.26)} = 2.99$, $p = 0.0599$) highlighting a faster implicit time in the resilient ($M = 32.29$, $SE = 1.01$) mice compared to the control ($M = 36.20$, $SE = 1.19$) mice ($p < .05$). The same observation is made between the susceptible ($M = 33.22$, $SE = 0.89$) and control mice even though it is not significant ($p = .0604$). Significance was determined using two-way mixed ANOVA with LSD multiple comparisons test. Symbols and bars represent the group average ± SEM; * $p < .05$, ** $p < .01$, *** $p < .001$; *n*-values are given in the method section.
